# Hybrid vesicle and reaction-diffusion modeling with STEPS

**DOI:** 10.1101/2023.05.08.539782

**Authors:** Iain Hepburn, Jules Lallouette, Weiliang Chen, Andrew R. Gallimore, Sarah Y. Nagasawa, Erik De Schutter

## Abstract

Vesicles carry out many essential functions within cells through the processes of endocytosis, exocytosis, and passive and active transport. This includes transporting and delivering molecules between different parts of the cell, and storing and releasing neurotransmitters in neurons. To date, computational simulation of these key biological players has been rather limited and has not advanced at the same pace as other aspects of cell modeling. As computational power advances and researchers want to add new realism to their models an important advance in the field of computational biology is to simulate vesicles in a realistic yet efficient manner. We describe a general vesicle modeling tool that has been designed for wide application to a variety of cell models, implemented within our voxel-based approach to modeling reaction-diffusion processes in realistic mesh reconstructions of cell tissue in our software STEPS. The implementation is validated in an extensive test suite, parallel performance demonstrated in a realistic synaptic bouton model, and example models are visualized in a Blender extension module.

## Introduction

Vesicles play many diverse and essential roles in cell biology. For example in neurons, synaptic vesicles uptake and store neurotransmitter, releasing the contents into extracellular space upon a presynaptic signal [1], and postsynaptic endocytosis and exocytosis of AMPAR-containing vesicles determine expression of AMPAR receptors in the cell surface [2, 3] via the vesicular endosomal pathway [4]. Therefore vesicles play central roles both in chemical communication between neurons and in determining the strength of synaptic contacts, two vital processes of brain function. Vesicles diverse in structure, molecular composition (even within functionally similar systems across anatomies, such as in the case of synaptic endocytosis [5]), and function play many more important roles throughout cell biology and are essential to life processes.

The field of computational biology has turned its attention to modeling these key cell biology players in recent years to varying degrees of biological detail. Many of these early applications are model-specific with a focus on one particular aspect of vesicle function that is being investigated in the study often alongside an experimental observation. Examples of this include an investigation of autophagy [6], vesicle mobility in the presynaptic space [7, 8], calcium-mediated release probability on docked synaptic vesicles [9], and the influence of cytoskeletal density on vesicle diffusivity [10]. A stochastic model of post-synaptic AMPAR trafficking by Tanaka and colleagues [11] is one of the most detailed vesicle models to date and includes many of the key features of the system such as endocytosis and AMPAR trafficking, but does have some simplifications and limitations such as drawing docking and exocytosis from random distributions instead of modeling full diffusion and with limited molecular complexity.

Whilst these early studies have paved a path to computational vesicle modeling, there is a need for a general tool or tools that have the flexibility to enable application to a variety of modeling systems. In addition, such a tool should also allow modeling of other features of the biological environment such as molecular reaction-diffusion processes and voltage-gated channels alongside the vesicle dynamics, in order to enable full modeling of the rich chemical processes behind important vesicle functions such as docking, priming and fusion.

In terms of designing a tool for such computational vesicle modeling, one possible approach is to model the forces that vesicles experience in the cytosolic environment explicitly by their interactions with proteins and other molecules in their environment (including water) to atomic detail, thus extending the field of molecular dynamics [12]. This approach has the benefit of high morphological and physical realism, however the drawback of such approaches is that the biological time that can be achieved in simulations is severely limited on current hardware due to the high intensity of computation, and often the experimental data of such atomic detail is absent [13]. Although coarse-grained approaches can increase the achievable biological time somewhat to the order of nanoseconds or microseconds [14], this still puts out of reach many models of interest in vesicle systems that may operate on a scale orders of magnitude higher such as synaptic vesicle endocytosis that operates on a timescale of seconds [15] within a synaptic vesicle cycle operating on a timescale of minutes [16].

In order to achieve these biological times, a good approach is to extend one of the current tools for subcellular modeling, bringing the many benefits of these established tools but with a vesicle-specific extension. STEPS is a voxel-based simulator that extends Gillespie’s Stochastic Simulation Algorithm (SSA) [17] to simulate diffusion within tetrahedral meshes that are able to capture complex biological boundaries accurately [18]. STEPS has been extended since its initial release to support voltage calculations on the tetrahedral mesh [19] and MPI-based parallel computation [20] via an operator-splitting framework [21]. The main benefit of SSA approaches is that they are fast and scalable [20], and capture reaction-diffusing kinetics accurately as long as certain geometrical criteria are met [18]. Although STEPS was originally designed with a focus on neural systems such as simulating molecular models of synaptic plasticity [22–24] or other dendritic spine molecular models [25], along with larger-scale dendritic systems [26–28], it has also been applied to other areas of biology such as to model viral RNA degradation and diffusion [29], astrocytic calcium signaling [30] and nanoparticle penetration in cancer research drug discovery [31]. The power of the MPI solver within STEPS was recently demonstrated by simulation of an entire Purkinje cell at molecular detail for seconds of biological time on a supercomputer [32].

In this study we describe how we take the many benefits of this parallel SSA-based simulator and extend it for modeling vesicles, which cannot realistically be represented by point-molecules within the regular SSA. Our aim to provide a flexible modeling tool that captures many important features of vesicles and their processes so that the tool can be applied to many different model systems. Vesicles are represented with enough realism to extract their key features such as mobility within cells and near membranes, their molecular interactions, and important features such as endocytosis and exocytosis, but by an approach that is capable of achieving biological times that are out of scope for molecular dynamics approaches. We describe our unique hybrid SSA-vesicle approach where vesicles and vesicle-related phenomena are given special treatment but with a firm focus on computational performance, and the benefits of the SSA are maintained for other chemical species and reaction-diffusion processes. We describe in some detail the implementation of our methods, and demonstrate biological plausibility by an extensive validation suite of all the new modeling components introduced in this paper. We introduce the parallel MPI-based implementation, demonstrate performance in a realistic synaptic bouton model, and visualize in a Blender extension module.

## Results

In this section we describe sequentially the features of vesicles that are modeled, and with results from test models by means of validating the method. We begin by qualitatively exploring vesicle mobility and crowding effects in the STEPS implementation. Later validations, however, are then generally run under well-mixed conditions (where crowding effects are negligible) in order to match to analytical solutions, although we should note in more complex models the reaction rates can be under-sampled or over-sampled due to effects of crowding.

Fig. 1 shows schematically the biological phenomena that STEPS simulates in the vesicle extension. Each of these features are described within the following sections.

**Fig. 1.**
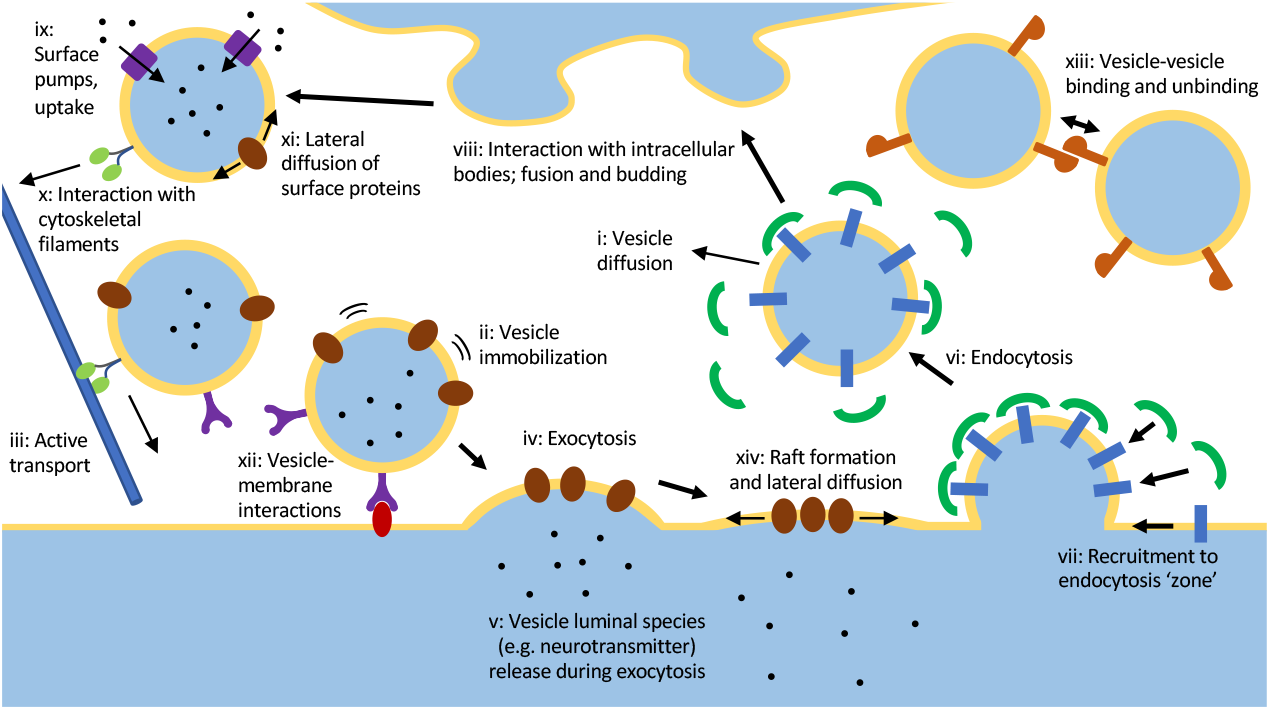
The full set of behaviors that STEPS implements for Vesicle and Raft modeling, described further in text. As such, models like the synaptic vesicle cycle including trafficking, docking, exocytosis and endocytosis are supported, as well as variations on those models such as kiss-and-run fusion and recycling. These vesicle models can exist alongside ‘regular’ reaction-diffusion and bioelectrical cellular models.

### Vesicle mobility: passive and active transport in the crowded cell environment, partial mobility in clusters, and tethering to the cell membrane

One of the most important features of vesicles is that they exhibit various forms of mobility, dependent on many factors such as location, function and cell activity [33]. They can, for example, be freely-diffusing within the crowded environment of the cytosol or other cell compartments (Fig. 1(i)), effectively immobile when docked to tethering molecules in the cell membrane (Fig. 1(ii)), or partially mobile when clustered to other vesicles [34] in functional ‘pools’ [35] (Fig. 1(xiii)). STEPS supports all of these situations. By default, vesicles freely diffuse within their cellular boundaries, although mobility is reduced in the presence of other vesicles or other internal cellular structures [8] thus capturing the important known features of cellular crowding [36]. To demonstrate this, first we investigated vesicle diffusion in the case where 0%, 20%, 40% and 60% of the cell volume is occupied, showing how the software captures the expected diffusion distance in the 0% case, and mobility reduces with increasing volume occupancy (Fig. 2a), as expected. In addition, we tested a mitochondria-based model introduced by Rothman et al [8]. In this model 28% of the volume is occupied by immobile mitochondria in the tetrahedral mesh, a further 25% by immobile vesicles, and 17% by mobile vesicles (the model is shown in the inset of Fig. 2b). We demonstrate that STEPS captures the expected features of this model in that the effective diffusion rate of the mobile vesicles is not constant in time; it approximates the Brownian-motion rate (0.06*μ*m^2^/s in this model) only on short time-scales and is reduced by steric interactions on longer timescales (Fig. 2b) approaching physiological values [8]. STEPS does not model the very fast timescale effects of hydrodynamics [8].

**Fig. 2.**
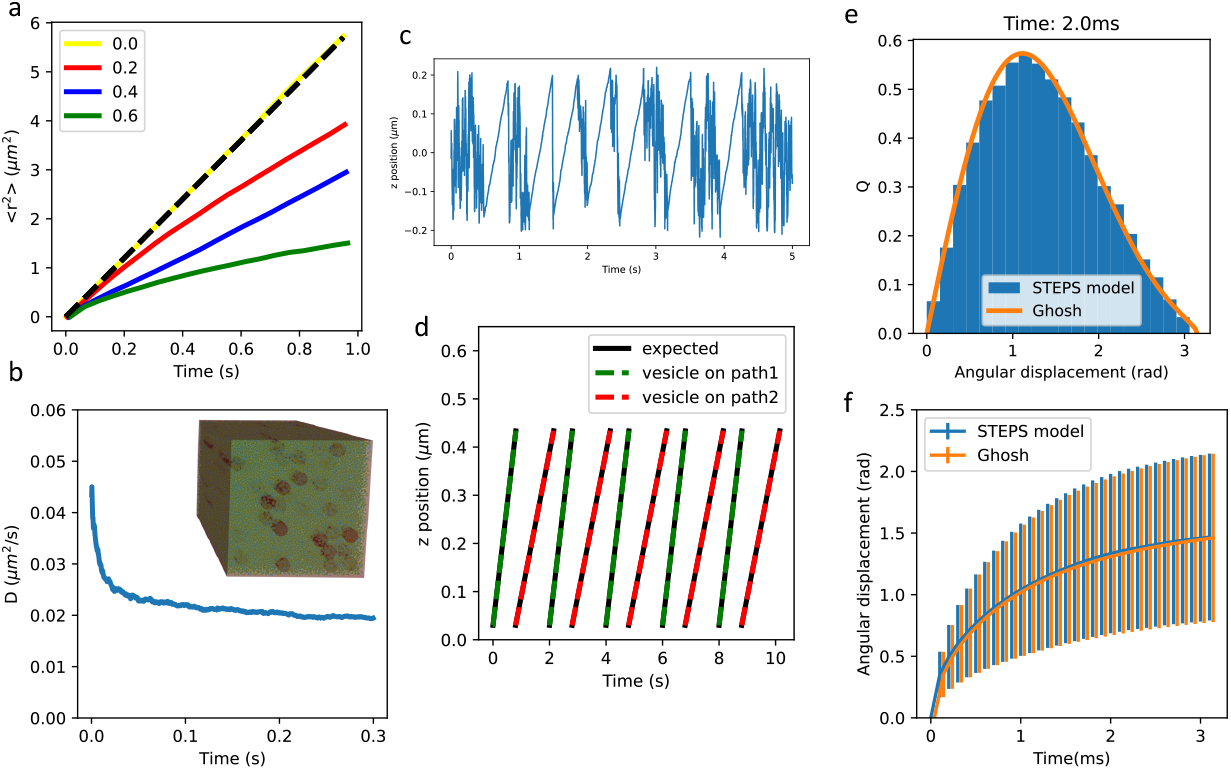
Validation of vesicle mobility, and mobility of molecules on vesicle surface. **a** Vesicle diffusion in a cytosolic volume with varying fractions of the volume occupied. Free diffusion (yellow line) is compared to expected 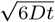 solution, and increasing cytosolic occupancy of 20%, 40% and 60% show increasingly reduced mobility, as expected. **b** The apparent diffusion coefficient of vesicles vs time in the STEPS implementation of the MFT model of Rotham et al [8] showing reduced apparent diffusion rate with time due to steric interactions with mitochondria and other vesicles, agreeing with the results from that study. **c** In an example model, vesicles exhibit brownian motion diffusion punctuated by periods of active transport due to interaction with actin filaments. **d** Validation of active transport, with vesicles maintaining the correct speeds upon interactions with ‘paths’ at 0.5um/s (green) and 0.3um/s (red). **e** The STEPS implementation (blue) of surface species angular displacement by the Ghosh algorithm [37] (orange) for an example diffusion step of 2ms for a surface species with diffusion rate on the vesicle surface of 1*μ*m^2^/s. **f** Mean and standard deviation of spherical surface diffusion over a range of diffusion timesteps. The Ghosh algorithm (orange) is compared to a surface diffusion model on a spherical mesh surface in STEPS (blue).

### Vesicle active transport

Active transport on structures such as microtubules and actin filaments (Fig. 1(iii,x)) is an essential feature of vesicle mobility, especially in neurons [38]. For example, synaptic vesicle trafficking along actin filaments delivers vesicles to the active zone [39], and post-synaptically recycling vesicles are actively transported to the spine surface to deliver AMPAR during LTP [40]. Vesicles are transported by motor proteins myosin on actin, and kinesin and dynein on microtubules [41]. A feature of the motor-protein ‘walk’ is static periods punctuated by sharp steps of fairly regular size to the next position along the structure [42–44]. The dwelltimes of which are near single-exponential or double-exponential [43, 45, 46], indicating one or sometimes two rate-limiting steps.

STEPS models active transport by a system of branched *Paths* representing cytoskeletal filaments that vesicles may interact with if they cross them in their environment. Vesicles travel along the path from the beginning point to the end point by a user-defined speed and stepsize (for example the stepsize of myosinV is *∼*36nm [41]). The default behavior is to sample dwelltimes from a single-exponential distribution, although a double-exponential may optionally be used. As well as active transport, paths may be useful in other modeling contexts such as vesicle binding to actin cytoskeleton in cluster formation [47]. In a simple model, the effect of paths can clearly be seen as periods of random diffusion interspersed by periods of regular movement from one end of the path to the other (Fig. 2(c)), mimicking the biology of periods of regular diffusion punctuated by periods of active transport [39, 48]. In addition we validate the implementation of paths in a model measuring vesicle displacement on path vs expected displacement, as demonstrated in Fig. 2(d)). The model vesicles undergo several periods of active transport along paths at the expected speeds (in this example with small stepsize to counteract stochastic effects for the purpose of validation). The implementation of this and all other validation models is described in further detail in the Supplementary Information.

### Vesicle surface molecule diffusion

Vesicles contain proteins in their surface which may have been present at the site of endocytosis (Fig. 1(vi)) and which can undergo various interactions such as phosphorylation upon interaction with the cytosolic environment. These vesicle surface molecules may play important roles such as neurotransmitter uptake ((Fig. 1(ix))), docking to cell membrane (Fig. 1(xii)) or binding to other vesicles (Fig. 1(xiii)). These molecules may be mobile in the vesicle surface, undergoing lateral diffusion (Fig. 1(xi)). This amounts to modeling diffusion on a spherical surface, by which STEPS applies the efficient ‘approximate propagator’ algorithm of Ghosh et al [37]. As described further in Supplementary Information, to test the STEPS implementation of the Ghosh algorithm we tested a regular diffusion model on a spherical surface mesh in STEPS and compared to the Ghosh algorithm (Fig. 2e shows one example of angular displacement after time 2ms of a surface molecule diffusing at 1μm^2^*/*s). The Ghosh algorithm compares accurately to the measured angular displacement distribution from our surface diffusion model (Fig. 2f) and is a reliable, fast algorithm for implementing diffusion of surface molecules on vesicles in STEPS.

### Exocytosis, endocytosis, fusion and budding

Exocytosis (Fig. 1(iv)) is an essential cell process by which luminal species inside vesicles can be secreted into the extracellular space (for example neurotransmitter release at chemical synapses) and membrane proteins may be inserted into the cell membrane. Both processes are vital to cell function. Vesicles are also able to fuse to internal membranes such as endosomes (Fig. 1(viii)), and both this and exocytosis are modeled in STEPS by the same computational framework within the SSA. STEPS is also able to model the complex chemical processes leading up to and resulting in exocytosis, such as those involved in docking, priming and fusion (Fig. 1(xii)) [49].

For synaptic vesicles it has been proposed that ‘kiss-and-run’ fusion [50], whereby a vesicle does not completely collapse upon neurotransmitter release but remains intact and is immediately endocytosed and recycled, is utilized in neurons due to the clear advantage of speed of recycling. Full-collapse fusion and kiss-and-run are proposed to coexist in many systems [51] with reports varying in the range of 5% and 80% for the vesicles undergoing kiss-and-run as opposed to full exocytosis [52]. Since the ability to model both modes of exocytosis is clearly desirable, STEPS supports modeling of both full-collapse fusion and kiss-and-run, with a user able to employ either or both modes within a model.

During kiss-and-run fusion, vesicles do not only preserve shape but can also partially preserve membrane protein and even luminal composition [51]. In a STEPS kiss-and-run fusion event, all luminal molecules such as neurotransmitter are released (Fig. 1(v)) but membrane-associated molecules may either be retained or effectively diffuse to membrane on a species-specific manner, at the modeler’s discretion. Within a full-collapse fusion exocytosis event in STEPS all luminal molecules are released into the extracellular space and all vesicle surface-bound species are transferred to the cell membrane (Fig. 1(iv, xiv)). As mentioned, fusion is also possible on internal structures (Fig. 1(viii)), enabling modeling of cycles such as the the post-synaptic AMPAR cycle by which AMPAR-containing vesicles fuse to endosome [4].

Fig. 3a provides a basic validation of exocytosis in STEPS within the SSA framework, with further details of this model given in the Supplementary Information. Fig. 3e visualizes a full-collapse fusion event in STEPS of one synaptic vesicle in the active zone, where release of neurotransmitter into the extracellular space and retention of the SNARE complex and vesicle surface-bound species in the cell membrane can be seen.

**Fig. 3.**
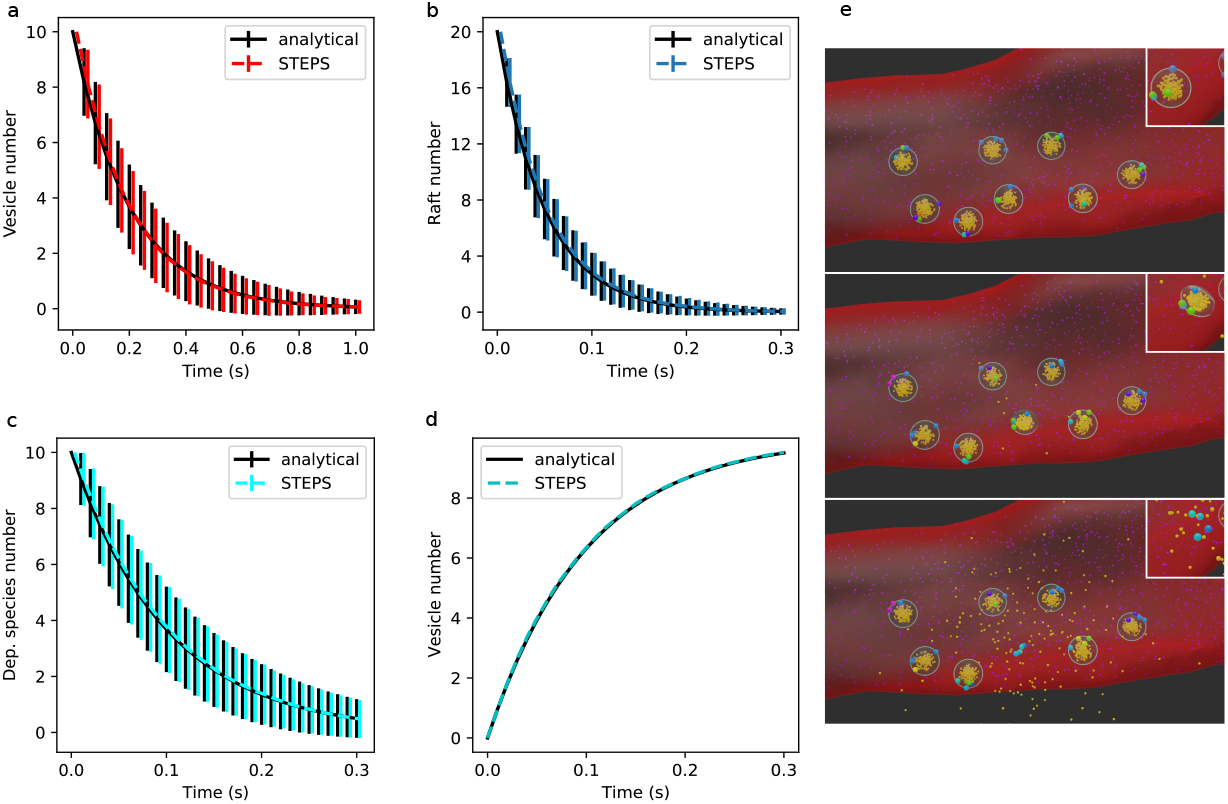
Validation of Endocytosis and Exocytosis in STEPS by basic test models where analytical comparisons can be made. **a** 10 vesicles undergo exocytosis at a rate of 5/s in a model where diffusion is not rate-limiting. Vesicle number decays exponentially, as expected. Mean and standard deviation of 1000 iterations are shown to validate correct implementation within the SSA. **b** Raft endocytosis is similarly validated as an exponential decay of 20 Rafts in the model at a rate of 20/s (mean and std of 1000 runs). **c**,**d** Endocytosis at rate 10/s is validated as exponential decay of a dependent species on membrane where 1 molecule is internalized per endocytosis event (**c**) and the increase in vesicle number due to endocytosis (**d**). **e** An exocytosis event visualized in Blender. Top panel: vesicles are docked in the active zone by SNARE complexes and calcium is released in the cytosol (purple). Middle panel: The vesicle in the center starts to undergo exocytosis. Bottom panel: Exocytosis is complete, neurotransmitter (dark yellow) is released into the extracellular space and the SNARE complex remains in the membrane for dismantling. The different colors of the SNARE complexes indicate different stages of priming, with the fully primed state on which exocytosis depends shown in light blue. The inset in each panel shows a close-up (from a different angle) of the vesicle undergoing exocytosis.

Endocytosis (Fig. 1(vi)) involves the creation of vesicles from an invaginated region of a cell membrane, and an analogous event termed ‘budding’ from internal structures such as liposomes [53] (Fig. 1(viii)), are modeled in STEPS as events within the SSA. These events can optionally be modeled with a species dependency, meaning that the event is only active and available for application within the SSA if a certain species signature is met, for example if a certain number of clathrin molecules have been recruited to the region [54] defined as an *Endocytosis Zone* within STEPS (Fig. 1(vii)). This region may be defined as a collection of surface triangles in the tetrahedral mesh, or as molecular population of a *Raft* (Rafts are described later in section Membrane Rafts: Formation, diffusion and interaction with their environment). Fig. 3b-d provides a basic validation of endocytosis in STEPS within the SSA framework when the endocytosis region is either based on rafts (Fig. 3b) or comprised of mesh surface triangles (Fig. 3c).

STEPS is capable of modeling the complex molecular machinery that generates endocytosis events, but does not yet support adaptive meshes and so the geometric changes are not fully modeled. Instead, when an endocytosis event is chosen to take place, the spherical space just inside the endocytosis region is reserved and a vesicle is formed in place. This vesicle will contain the molecular makeup of the zone in which endocytosis occurred in its membrane, to mimic the molecular biology (Fig. 1(vi)). Fig. 3d validates the expected production of vesicles during repeated endocytosis events. As dependent species are consumed the rate of production of vesicles can be seen to decrease in this validation model.

### Vesicle surface protein and vesicle internal molecule transport, and interaction with environment

Vesicles contain proteins embedded in their membrane which allow them to interact with their environment through protein-protein or other interactions (Fig. 1(ix, xii)), or even interact with each other (Fig. 1(xiii)). An important example of such interactions are those that allow synaptic vesicles to go through the process of docking and priming, which is controlled by interactions between vesicle proteins and proteins embedded in the cell membrane [55]. Further, primed vesicles then sense calcium released into the cytosol to undergo the process of fusion and neurotransmitter release through a cascade of chemical interactions [49, 56].

STEPS supports all of these situations by addition of a new type of reaction termed a *Vesicle Surface Reaction* by which reactants may exist in a number of regions: they may be compartment-based (e.g. cytosolic) molecules, molecules in the vesicle surface, or molecules embedded in the cell membrane or other membranes. A reaction between a molecule in the vesicle surface and the cell membrane can be used to model docking (Fig. 1(xii)), for example. In addition, reaction products may appear internally within the vesicle to model e.g. neurotransmitter pumps and filling (Fig. 1(ix)). These reactions are then solved stochastically as regular SSA reactions. This allows modeling of the rich interactions that vesicles may undergo with their environment and allows, for example, the entire synaptic vesicle cycle to be modeled to full molecular detail [57]. To faithfully model docking reactions specifically, optionally a maximum distance may be defined so that the reaction will only occur within a specified distance of the membrane surface, and vesicles may be immobilized when effectively docked close to the membrane (Fig. 1(ii)). The reverse reaction can also be modeled where vesicles may become undocked and mobilized, which is important if modeling kiss-and-run fusion.

Fig. 4 demonstrates our validation of the vesicle surface reactions by way of interaction with vesicle surface molecules and cytosolic species. We extend our regular validations [18] for these interactions, and demonstrate very close agreement in four different models, namely first order irreversible (Fig. 4a) and reversible (Fig. 4b) reactions, and second-order irreversible (Fig. 4c) and reversible (Fig. 4d) reactions, in every model demonstrating very close agreement to the expected solution analytically. Supplementary Information contains further details of the model setup and parameters used.

**Fig. 4.**
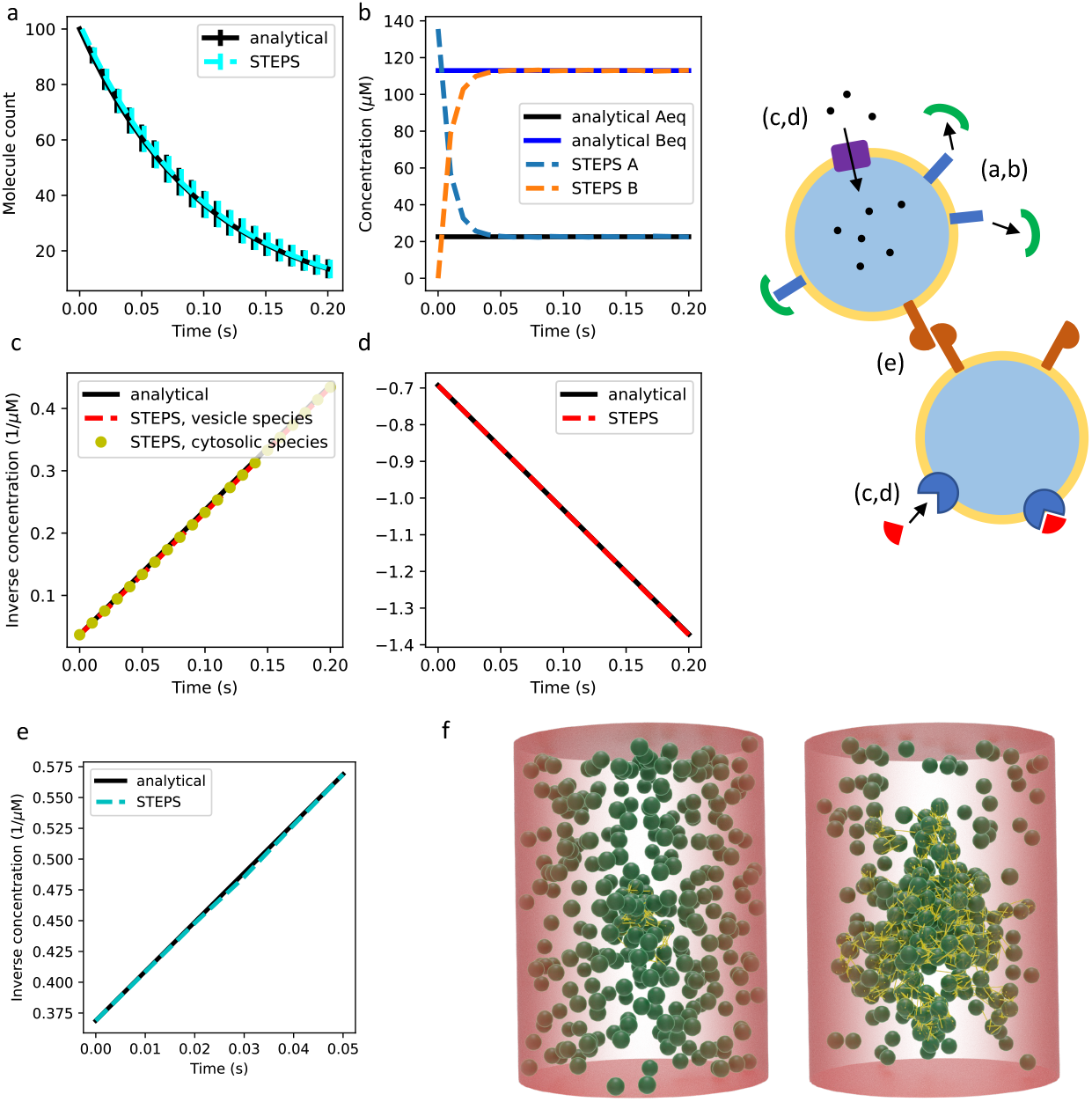
Validation of vesicle surface molecule interactions. As described further in text and Supplementary Information, **a** and **b** validate first order surface interactions whereas **c** and **d** validate second-order interactions such as interactions with cytosolic proteins and surface pump action. **e** Validates vesicle binding interaction (vesicles binding to each other) which can be visualized in **f** as Blender output of a clustering model showing various stages of dynamic cluster formation (the synapsin dimer proteins that bind vesicles to each other are shown in yellow). The cluster can form or disperse based on cellular activity in the model.

### Vesicle interactions with each other

A known behavior of vesicles is their ability to form functional clusters or ‘pools’ [35, 58] in which mobility is strongly reduced [59]. It has been proposed that such clusters are formed by vesicles tethering to each other for example in the case of synaptic vesicles by synapsin interactions [47, 60]. STEPS models this phenomenon by allowing vesicles to loosely bind to each other through interactions between their surface proteins. A special type of reaction termed a *Vesicle Binding* event models this, alongside the reverse process *Vesicle Unbinding* (Fig. 1(xiii)).

As vesicles bind in STEPS they form a computationally distinct complex termed a *Link Species* on their surface (Fig. 4f). Link Species have the property of length, which is by a user-defined upper and lower boundary, and the Vesicle Binding event can only occur if and when the resulting Link Species formed would be within its permissible bounds. Link Species allow some mobility of linked vesicles as long as any movement results in the Link Species remaining within its bounds. In addition, Link Species can undergo all the usual chemical interactions such as interaction with cytosolic or vesicle-bound molecules, and diffusion on the vesicle surface. Through this behavior, vesicles are therefore able to form clusters in which they exhibit loose, partial mobility, phenomenologically similar to their low mobility in pools [59]. Thus vesicle clustering and pool formation such as the classical reserve pool, recycling pool and readily releasable pool [35] can be faithfully modeled in STEPS, along with their interactions such as activation by calcium, and the process can be modeled to high molecular detail.

We provide a validation model of the Vesicle Binding reaction in STEPS. Since this follows a second order [A]+[A] type reaction there is a known analytical solution to which STEPS output can be compared. Fig. 4e shows close agreement between the STEPS model and analytically expected behavior of this model. In addition, Fig. 4f demonstrates cluster formation in which synapsin dimers bind vesicles to each other, forming a fluid cluster [34].

### Membrane Rafts: Formation, diffusion and interaction with their environment

Nanoscale domains of clustered proteins, lipids and other molecules in cell membranes play many important roles in signaling and trafficking [61]. Referred to as ‘Lipid Rafts’, these important membrane subdomains span sizes from *∼*50nm to hundreds of nanometers, are dynamic in their molecular composition, and may be mobile and diffuse freely in the membrane or become bound to cytoskeleton and immobile [62]. Somewhat analogously to vesicles within cytosol and other cell compartments, rafts cannot be modelled as regular volumeless molecules within SSA methods and require special treatment.

We implement rafts in STEPS as regions that occupy an exclusive, fixed radius within cell membranes. To mimic the biology, rafts may be mobile by a user-defined diffusion rate within cell surfaces (Fig. 1(xiv)) or immobile. They can undergo all the usual chemical interactions with their environment including with cytoplasmic molecules and surface-bound molecules and, in addition, rafts can be created upon exocytosis of a vesicle (as proposed for example in [63]) or form an endocytic zone from which a vesicle will be produced, sharing the surface molecular composition with the exocytosed or endocytosed vesicle (Fig. 1(xiv, vi)).

Since rafts are associated with a triangular meshed surface in a STEPS simulation, we apply the same diffusion rule for rafts as for regular SSA surface species. Fig. 5a validates this implementation in a diffusion model where crowding effects are minimal, although it should be noted that diffusion rates can be affected by crowding if raft density is high. In addition, and similarly to our validation of vesicle surface reactions, we validate our implementation of raft surface reactions by models of first-order irreversible (Fig. 5b) and reversible (Fig. 5c) reactions, second-order irreversible (Fig. 5e) and reversible (Fig. 5f) reactions. As a useful modeling feature, it is also possible in STEPS to model formation of rafts when a particular species signature is met, and conversely dissociation and dispersion can be modeled if a raft molecule composition goes below a certain threshold. Fig. 5d validates these implementations in a simple model that includes both of these phenomena. Further details of all of these models can be found in the Supplementary Information.

**Fig. 5.**
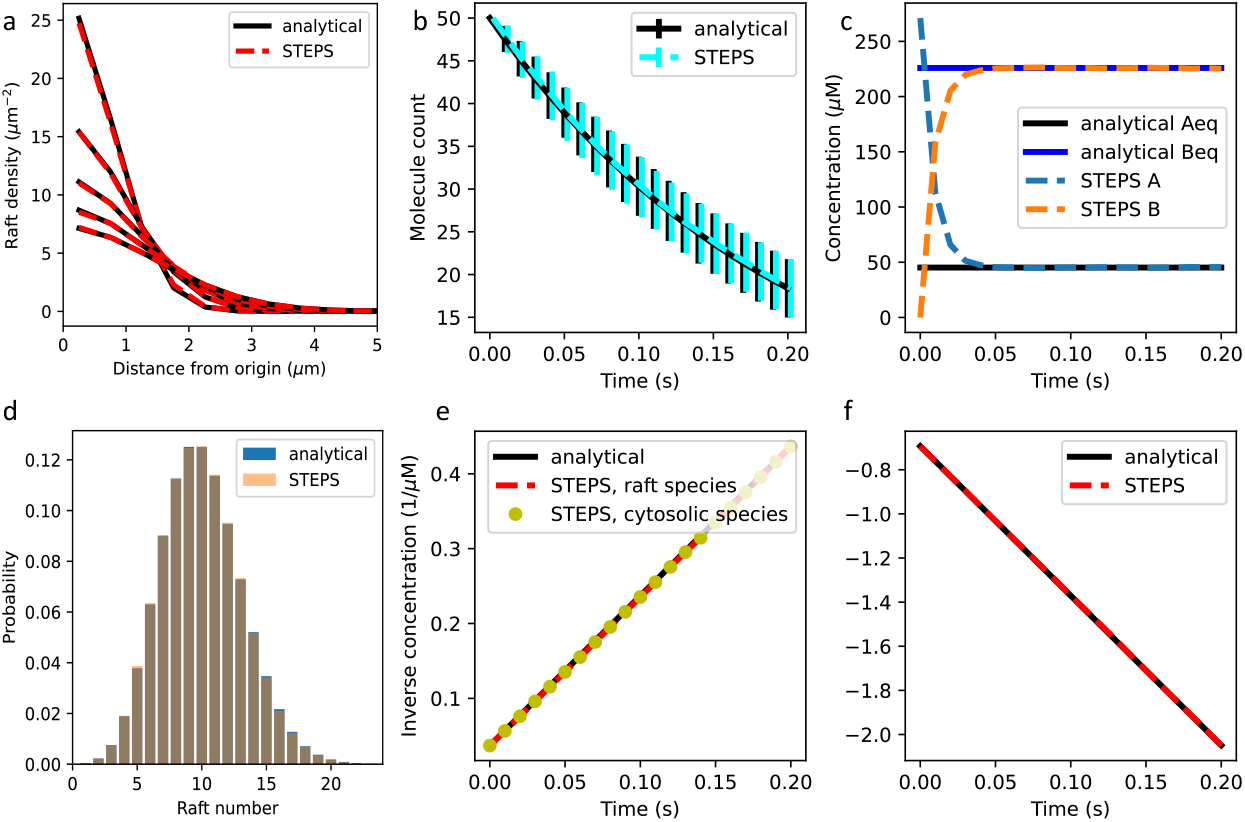
Validation of rafts and their interactions, described further in text. **a** Raft diffusion on membranes under no-crowding conditions. **b** First-order irreversible and **c** first-order reversible interactions. **d** Combined raft generation and raft dissociation interaction model validates the SSA implementation of these phenomena. **e** Second order irreversible interactions and **f** second-order reversible interactions with membrane-bound molecules and cytosolic molecules. All models are compared to analytical solutions.

### Computational performance in parallel

As described in further detail in MPI implementation, STEPS implements a parallel splitting algorithm for the vesicle solver, since an original serial implementation was found to be very limited in applications due to all the extra computation involved in supporting vesicle and raft modeling. We tested our MPI-based solution on a realistic and detailed synaptic bouton model further described in [57] and visualized in Fig. 6d.

**Fig. 6.**
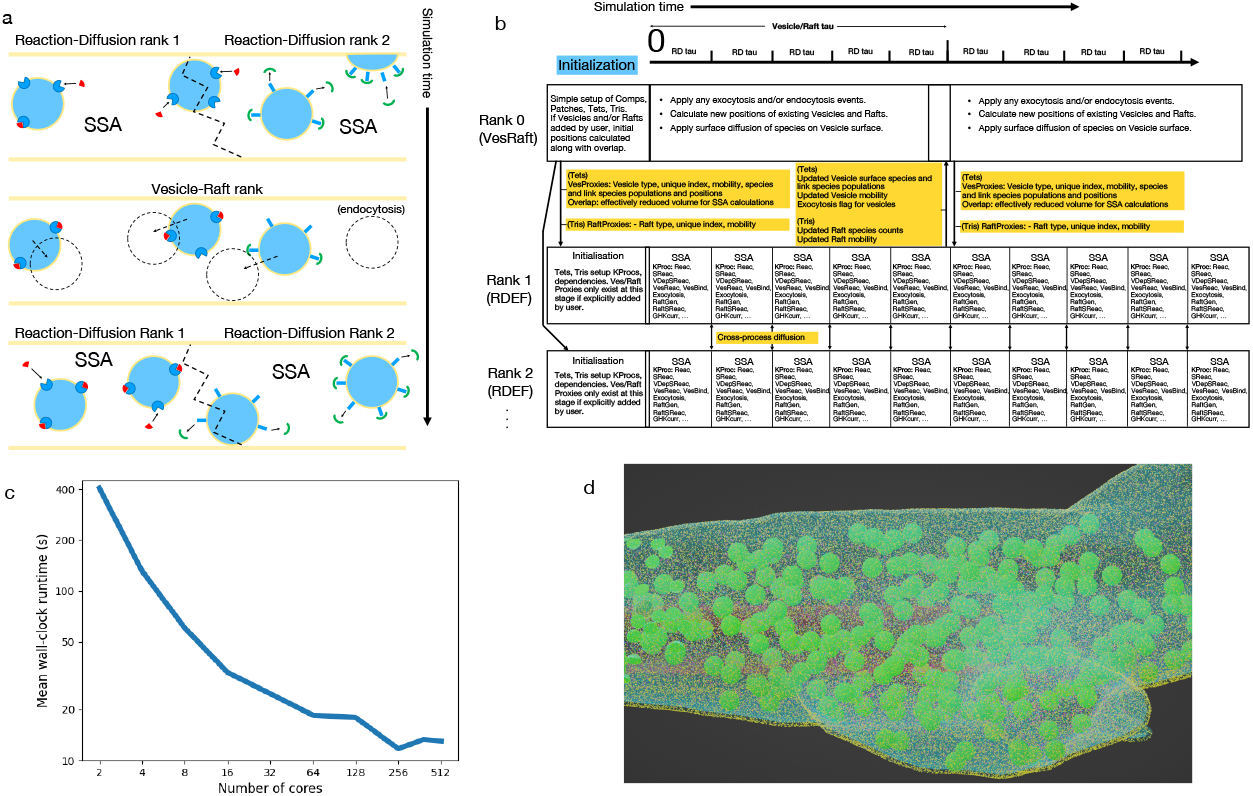
The MPI implementation and Blender extension module. **a** Schematic of the MPI ‘splitting’ solution, in which n-1 ‘RDEF’ ranks carry out reaction-diffusion and voltage calculations on partitioned mesh space, and on a slower clock the ‘VesRaft’ rank (rank 0) calculates vesicle and raft diffusion and some other vesicle-related processes over all space, communicating new positions to the RDEF ranks. **b** Detailed description of the parallel algorithm timeline. **c** Example parallel performance of a realistic synaptic bouton model of [57]. **d** Screenshot of the Blender extension model applied to the same synaptic bouton model.

In this stringent test of our initial MPI implementation, which is far from an idealistic scenario but instead a realistic biological scenario, we achieve a good reduction in simulation wall-clock time from 404s on 2 cores (the lowest number of cores possible in our implementation) to 11.7s on 256 cores (Fig. 6c) per 1ms of biological time. Although there is no further improvement in runtime with more cores beyond this point, this speedup is sufficient to bring systems that operate on timescales of seconds or minutes such as the synaptic vesicle cycle into scope. If cluster runtime constraints are a concern, checkpointing further extends the biological time that is achievable. Future improvements to MPI performance may be possible by parallelizing the vesicle routines and with further optimization to the MPI communication schemes.

## Discussion

In this study we presented a new modeling tool that supports realistic simulation of vesicles and their many important biological functions such as uptake, transport and secretion, as well as lipid rafts. To our knowledge, this is the first approach to successfully combine such modeling in a hybrid framework with spatial SSA modeling, enabling detailed chemical modeling of the processes involved. Every stage of development was carefully tested and validated, with many of the validations described in this paper, and implemented in ‘real world’ biological scenarios. In one example which we show in Fig. 6d and described further in [57], a detailed pre-synaptic bouton model simulated the synaptic vesicle cycle to full molecular detail.

The software was implemented in a novel MPI framework that took into account the unique challenges of modeling volumetric objects within the partitioned spatial SSA framework. Although this is a difficult system to parallelize, by taking advantage of the fact that vesicles diffuse much slower than smaller molecules we were able to sufficiently improve performance by orders of magnitude compared to a serial implementation, putting in reach biological systems operating on timescales of seconds or minutes. Such timescales are unreachable in molecular dynamic approaches.

We have developed this as a general tool that has the flexibility to enable application to a variety of modeling systems with realistically modeling of vesicles and lipid rafts and with the power to reach size and time scales of interest. The software is free for anyone to use under the GNU General Public License v3.

## Methods

### Vesicle diffusion

Vesicles are approximated in STEPS as spherical volumes as obviously the most natural choice of the available geometrical primitives. This volumetric modeling of vesicles is a clear difference to the effectively point-particle SSA species in STEPS used to model smaller molecules such as ions, ion-channels and enzymes. Cellular vesicles vary in size; for example in neurons synaptic vesicles are approximately 40nm in diameter and can transport a relatively small amount of neurotransmitter and surface proteins but can release quickly, whereas dense-core vesicles that transport neuropeptides are larger at 80-200nm and operate more slowly [64]. A STEPS user specifies vesicle diameter that vesicles of this type will then assume during the simulation as effectively a sphere that operates within the realms of the tetrahedral mesh environment. A STEPS user is not limited in how many different types of vesicles they may define. The user must then also define a fixed timestep on which to update vesicle positions by diffusion, which should obviously be kept small enough that diffusive steps are reasonably short. For example, for a diffusion coefficient of mobile synaptic vesicles of 0.11μm^2^*/*s [65], a timestep of 1ms will give a diffusion distance of *∼*10nm and so is a reasonable choice for most modeling scenarios.

On these regular timesteps *δt*, a diffusing vesicle’s next position is chosen by sampling the x, y, z displacement separately:

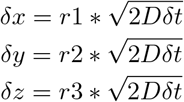

where r1, r2, r3 are random numbers generated on the standard normal distribution. The new position is then tested to see if it is a valid position or not. For example, the vesicle must not overlap any other vesicles nor must it cross a boundary such as the mesh surface. To optimize the search of other vesicles, only vesicles known to overlap the tested tetrahedrons are checked. To calculate whether a vesicle crossed a compartmental boundary requires knowledge of the sphere-tetrahedron overlap of the new position.

### Sphere-Tetrahedron overlap

STEPS uses an external library to determine the sphere-tetrahedron overlap [66] so that the set of tetrahedrons that a vesicle overlaps can be found. For application of this algorithm, it is essential to avoid testing all tetrahedrons in the entire mesh due to runtime concerns. If a vesicle is diffusing within a short distance inside its own diameter, the set of tetrahedrons previously overlapped are taken as the starting point for the overlap search because there is guaranteed overlap in this set. If a vesicle is being newly introduced into the mesh environment (for example by endocytosis or by a modeler API call) a walking algorithm is employed to find the starting point. A breadth-first search is then used to test overlap of tetrahedrons, stopping if no overlap can be found within a layer or overlap reaches 100% of the vesicle volume (which means that the new position is good). Overlap under 100% means that a position being tested is outside of the permitted environment (i.e outside of compartmental boundaries).

### Vesicle-mesh overlap effects on molecular reaction and diffusion

To approximate the effect of vesicle presence on the mobility of other molecules (modeled within the operator-splitting framework [21]), we apply a correction to the finite-volume diffusion rate [18] of molecules proportional to the volume occupancy. The modified diffusion rate from tetrahedron *k* to tetrahedron *l* of species *i* is:

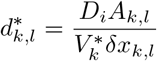

where *D*_*i*_ is the macroscopic diffusion coefficient of species i, *A*_*k,l*_ is the face area between tetrahedrons k and l, 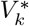 is the reduced volume by vesicle occupancy, and *δx*_*k,l*_ is the barycenter-barycenter distance. This reduced volume effect intuitively reduces the dwelltime for a molecule inside a tetrahedron proportional to the reduced volume, whilst also ensuring SSA species cannot occupy a fully overlapped tetrahedron and so do not occupy the same space as a vesicle. Note that mathematically the diffusion rate will approach infinity as the reduced tetrahedral volume approaches zero (meaning complete coverage by a vesicle) and is effectively replaced by a high rate for diffusion to a neighbor within the operator-splitting framework if this situation occurs. In practical tests, however, usually full coverage does not occur (and won’t occur unless tetrahedrons are very small) and in addition molecules are expected to evacuate tetrahedrons before full overlap occurs. Diffusion to a fully-overlapped tetrahedron is not permitted.

We tested the effect on regular diffusing SSA species in a model with 10% volume occupancy by vesicles. In all cases, the tetrahedrons captured the binomial distribution of molecules with low error, as shown in Supplementary Information. In all cases, the error in the fit to the binomial distribution was less than 0.5%. This means that no spatial bias is introduced on the SSA species distributions by the modified diffusion rates.

STEPS also applies a volume correction to SSA reaction rates in order to capture effects of a reduced volume. For example, the rate, *c*, of a particular second-order reaction taking place in tetrahedron *j* is:

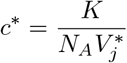

where *K* is the macroscopic reaction rate, *N*_*A*_ is Avogadro’s number and 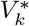 is the reduced volume by vesicle overlap. In this sense, the volume that vesicles occupy is not regarded as part of the ‘well-mixed’ volume of a tetrahedron. This can be expected to accentuate small-volume errors on reaction rates [67] if overlap is high. However, in practice, overlap is usually rather small so that this effect is also small or negligible because the effective volume is still usually within the accurate size window [18]. It should be noted that all validations presented in this paper, such as the vesicle surface molecule interactions presented in Vesicle surface protein and vesicle internal molecule framework. First order reactions are unaffected.

### MPI implementation

We previously parallelized the reaction-diffusion and voltage solution in STEPS [20], yet the complex additional vesicle-related functionality requires a customized solution that builds on this scheme. Our approach is to implement a splitting method whereby rank 0 (which we term the ‘VesRaft’ rank) carries out vesicle-related computations such as vesicle and raft diffusion, diffusion of vesicle surface species, and application of endocytosis and exocytosis (Fig. 6a,b), and the other available processes (which we term the ‘RDEF’ ranks) carry out the regular SSA calculations based on our previous method, which now however include many new phenomena such as reactions on the vesicle and raft surfaces, fusion and so on (Fig. 6a,b). This is achieved by effectively partitioning vesicles and rafts on the RDEF cores into vesicle and raft ‘proxies’ (Fig. 6a). This is essential to maintain the partitioned-SSA method that is known to scale well in realistic reaction-diffusion models [20], with many vesicle-related phenomena operating within the SSA in parallel. Of course this requires frequent communication between the VesRaft core and the RDEF cores (Fig. 6b).

The new solver inherits the MPI communication scheme for reaction-diffusion and the voltage solutions in the previous operator-splitting implementation [20], while adding communications of complex data structures between the VesRaft rank and the RDEF ranks (Fig. 6b). These two types of solver contain different local data, thus extra synchronization in data inquiry functions are also needed. These new challenges are addressed by the implementation of customized MPI datatype encapsulation and conditional MPI operation templates. To simplify and reduce the usage of MPI communication of multiple data entries in simulation objects (vesicles for example), data entries in object classes are firstly defined and encapsulated as custom datatypes using MPI’s user-defined datatype mechanism. Such encapsulation, together with our STL (Standard Template Library) container based MPI broadcasting and gathering templates, allows us to pack objects with multiple primitive data types as a single vector and synchronize using minimal MPI calls. This helps to reduce the latency due to frequent synchronization of small size data entries and improves the simulation efficiency. Data output in the VesRaft rank is often irrelevant to RDEF ranks, and vice versa, therefore global synchronization is not necessary, particular for vesicle-related data. We allow the user to switch communication mode for data inquiry functions by setting two condition flags in the solver: first, whether global output synchronization should be enabled; and second, if global output synchronization is disabled, which rank should receive the actual output. Note that network communication may still occur if the acquired data is not available in the output rank solver. In practice, we enable global synchronization by default, but disable it and set the VesRaft rank as output rank for heavy vesicle manipulation routines. Internally, this functionality is accomplished by a series of conditional MPI communication and data reduction templates, which automatically identify the data availability and perform the necessary data synchronization according to the conditions.

### Visualization with Blender

With the release of STEPS 4.1, the possibility to save simulation data to the HDF5 [68] format was added. This data can then be loaded into scientific data visualization software such as Paraview [69]. Although well suited to the visualization of purely mesh-based data, this approach was not ideal for the hybrid SSA-vesicle simulations described in this paper. We therefore developed a separate `stepsblender` python package that can read simulation and mesh data automatically saved to an HDF5 file and visualize it in the Blender 3D computer graphics software [70]. Simulations can be visualized interactively or rendered as movies directly from Blender. The visualization includes vesicles, rafts, link species, regular species, as well as compartments, patches, vesicle paths, and endocytic zones. Fig. 6d shows an example still shot from the model described in [57] and output from the Blender extension module is also shown in Fig. 2b, Fig. 3e and Fig. 4f.

This new python package contains a command line tool to import the data into Blender as well as python modules that can be imported in python scripts to further customize the appearance of model objects. To avoid memory issues in Blender, only the data corresponding to the currently visualized timestep is loaded into Blender. The data loading from HDF5 file is handled in a separate process, potentially running on a different machine, allowing the visualization of data from a computing cluster without downloading the full HDF5 files.

## Code availability

STEPS is available from https://steps.sourceforge.net

## Acknowledgments

We thank Tristan Carel of the Blue Brain Project, École Polytechnique Fédérale de Lausanne for his contributions to the vesicle code.

## Author information

### Contributions

EDS conceptualized and led the study. IH led software development. JL and WC contributed to software development and IH, JL and WC did pre-release testing and debugging. IH designed and carried out the validation studies. JL wrote the Blender visualization module. AG and SYN designed features of the software and tested the implementations. IH, JL, WC and EDS wrote the manuscript. All authors read and approved the final version.

