## Supplementary Information for "Hybrid vesicle and reaction-diffusion modeling with STEPS"

### 1) Vesicle diffusion

A single vesicle of diameter 50nm was injected into the centre of a tetrahedral mesh of a 10μm radius sphere. The vesicle was given a diffusion rate of 1 $\mu m^2/s$ and the simulation was run for 1s, with the distance from the origin measured at 0.05s intervals. This simulation was repeated 10,000 times with the mean taken for the data shown in Figure 2a (yellow). For the cytosolic occupancy simulations, tetrahedrons representing a total of 20%, 40% and 60% of the overall simulation volume were randomly removed from the simulation effectively creating cytosolic obstructions, for the same simulation protocol as above.

### 2) Mossy Fibre Terminal simulations: how steric interactions affect vesicle mobility

To mimic the simulation conditions of Rothman et al [1], a tetrahedral mesh of the Mossy Fibre Terminal volume of cubic boundaries 3μm x 3μm x 7μm was created with 565,958 tetrahedrons. Cylindrical mitochondria of 0.3μm diameter and 2.25μm length on the z-axis were created at random positions within the volume up to a total volume occupancy of 28%, numbering 110 in total, creating fixed obstructions within the cytosolic volume.
For the simulation, two types of vesicles were used – mobile vesicles, making up 17% of the overall MFT volume, and immobile vesicles making up 25%. The mobile vesicles were given a diffusion coefficient of 0.06$\mu m^2/s$ and all vesicles had a diameter of 49nm. The initial state of the simulation is shown in the inset of Figure 2b. The simulation was run for 0.3s and the mean square displacement of mobile vesicles starting within a 1μm radius of the mesh centre was recorded. This was divided by the time vector and 6 to give an apparent diffusion rate vs time (because $\langle R^2 \rangle = 6Dt$), as shown in Figure 2b.

### 3) Active transport

Active transport is simulated in STEPS on simulation objects called 'Paths' that represent cytoskeletal filaments such as actin or microtubules and may be branched. A user-defined stepsize and mean speed for vesicle active transport along these paths is then used to simulate fixed-sized steps from start to end. The time period until the next step ('dwelltime') is taken either from a single-exponential or double-exponential which is calculated from the given parameters.

Figure S1 shows the sampled 1-step (single-exponential) and 2-step (double-exponential) distributions, and recorded distributions from a STEPS simulation. In this simulation, the vesicle travels along the path at a mean speed of 0.8μm/s with a stepsize of 36nm, giving a mean dwelltime of 0.045s.

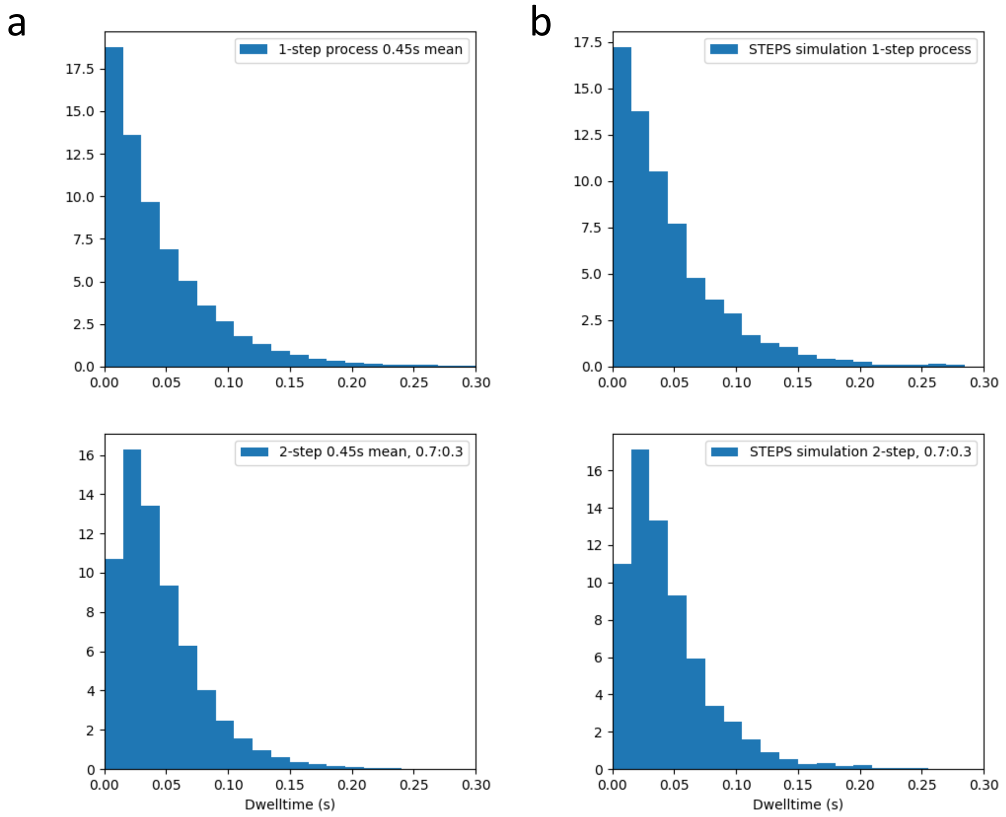


*Figure S1. a) Single-exponential (top panel) and double-exponential in 0.7:0.3 ratio (bottom panel) dwelltime for a vesicle undergoing active transport on a path in STEPS, and b) the recorded dwelltimes taken from a STEPS simulation, validating the implementation.*

### 4) Free diffusion with periods of active transport

In the simple model shown in Figure 2c, a single vesicle of 50nm diameter was placed randomly inside a tetrahedral mesh of a 0.5μm diameter sphere centred on the origin, and allowed to freely diffuse at $1\mu m^2/s$. A single path of length 0.35μm was placed inside the volume on the z-axis and the vesicle specified to undergo active transport on the path with a speed of 1μm/s. The z-position of the vesicle vs time is shown in Figure 2c in order to clearly see the interaction with the path on the z-axis.

### 5) Active transport validation

We created a model including two distinct paths with vesicles interacting by different speeds on each path in order to validate the active transport implementation in STEPS. The vesicles moved with mean speed of 0.5μm/s on the $1^{st}$ path and 0.3μm/s on the $2^{nd}$ path. Vesicles will optionally only interact with a path if a given species dependency is met, and this was utilised in the model to ensure vesicles interacted with the expected path at the expected time. Vesicles were injected at the start of the paths at 2s intermittent time intervals starting at 0s on the 1st path and 0.8s on the 2nd path. Both paths were on the z-axis, and the z-position of the vesicle was recorded and compared to the expected position based on the speed as shown in Figure 2d.

### 6) Vesicle surface molecule diffusion

To perform our own validation of the approximate propagator algorithm of Ghosh et al [2] for surface diffusion on vesicles, as implemented in STEPS, we designed a model in which surface diffusion on a spherical mesh could be compared directly to output from the algorithm. We created a tetrahedral mesh of a sphere of radius 50nm and simulated diffusion on the surface in STEPS. The surface diffusing species was given a diffusion coefficient of $1\mu m^2/s$ and the model was simulated for 3ms over 100,000 iterations. The angular displacement distribution was then compared to the Ghosh algorithm at 2ms, as shown in Figure 2e, and the angular displacement mean and standard deviation at 0.1ms intervals were compared over the whole 3ms simulation as shown in Figure 2f. Supplementary Figure S2 shows the full distribution also at 1ms (a) and 3ms (b), and compares the Ghosh algorithm and STEPS implementation to naïve planar diffusion (c).

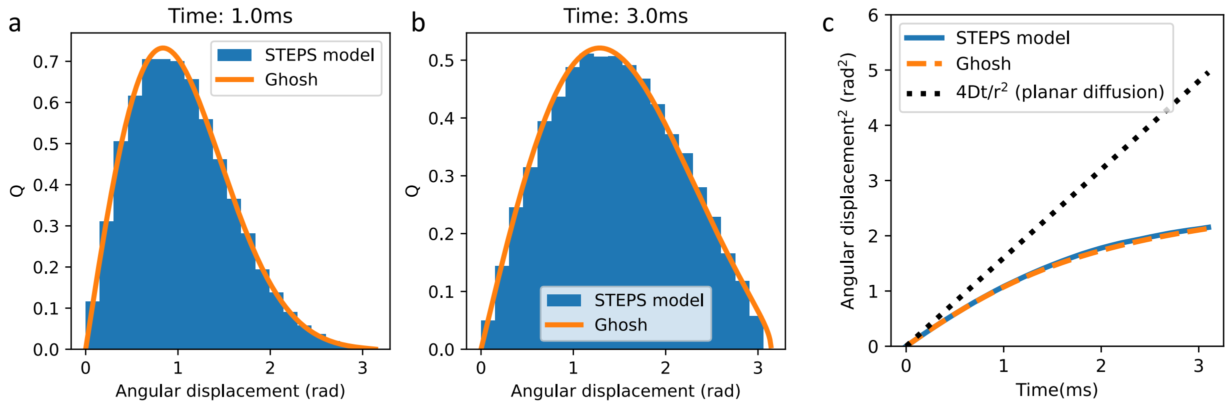


*Figure S2. The STEPS model of surface diffusion on a sphere compared to the Ghosh approximate propagator algorithm. a) angular displacement distribution at 1ms and b) 2ms. c) The STEPS surface diffusion model (blue) and Ghosh algorithm (orange) shown as mean output and compared to simple planar diffusion.*

### 7) Exocytosis

To validate the implementation of exocytosis with the SSA, we created a model in which 10 vesicles of 40nm diameter were free to diffuse around a spherical volume of 0.5$\mu$m diameter at a diffusion rate of $1\mu m^2/s$. Exocytosis could then occur anywhere on the membrane at a rate, k, of 5/s. Under these conditions, diffusion is not rate limiting and with exocytosis implemented as a first-order reaction the vesicle population should follow the expected exponential decay:

$$N(t) = N_0 e^{-kt} \quad (1)$$

And the standard deviation also has a known analytical solution:

$$\sigma(t) = \sqrt{N_0 e^{-kt}(1 - e^{-kt})} \quad (2)$$

We simulated the exocytosis model for 0.2s and compared the mean and standard deviation of 1000 runs to equations 1 and 2, as shown in Figure 3a.

### 8) Endocytosis

We used two models to validate Endocytosis in STEPS. In the first model, endocytosis occurred from membrane Rafts. On the surface of a mesh of a sphere of 0.5μm diameter, 20 immobile rafts were randomly placed and specified to undergo endocytosis at a rate of 20/s. The resulting raft number on the surface is then expected to follow an exponential decay following equations 1 and 2, and 1000 iterations of the model were run in order to make the comparison to the analytical solution, as shown in Figure 3b.

In the second model the alternative mode of endocytosis was tested, occurring when a certain species signature is met within surface mesh triangles. In this simple test model, a dependency of just 1 molecule was used with endocytosis occurring at a rate of 10/s, and with 10 molecules randomly placed on the mesh surface. Since the dependent species is absorbed during endocytosis and becomes located on the surface of the endocytosed vesicle, the dependent species population in the mesh surface is also expected to follow an exponential decay following equations 1 and 2, with the mean and standard deviation of 1000 iterations shown in in Figure 3c. For this model, the vesicle number is also shown in Figure 3d, which is expected to rise as:

$$N(t) = N_0 + M_0(1 - e^{-kt}) \quad (3)$$

where N is the vesicle number (in this model $N_0 = 0$, the vesicle number is 0 at time 0) and $M_0$ is the initial dependent species number.

### 9) Vesicle surface molecule interactions

To validate the implementation of vesicle interactions with their environment through molecules in their surface, we tested 4 different types of reactions occurring on 10 vesicles of diameter 40nm freely diffusing around a spherical volume of 0.5μm diameter at a rate of 1μm$^2$/s:

- A first-order irreversible reaction following equations 1 and 2 and shown in Figure 4a. For this model, a total of 100 molecules were injected (10 per vesicle) and allowed to decay at a rate of 10/s.

- A first-order reversible reaction: $A \underset{k_b}{\overset{k_f}{\rightleftarrows}} B$

If $[B]_0$=0 for simplicity, the expected equilibrium concentrations are:

$$[A]_{eq} = \frac{k_b[A]_0}{k_f+k_b} \quad (4)$$

$$[B]_{eq} = [A]_0 - [A]_{eq} \quad (5)$$

Where [ ] indicates cytosolic concentrations. The reversible reaction was implemented on the vesicle surface with 500 molecules of A per vesicle surface (which is quite high to reduce the effects of noise) giving $[A]_0$ = 135.6μM with $[B]_0$ = 0μM, and with $k_f$=100/s and $k_b$=20/s. The mean cytosolic concentrations over 10 runs are shown in Figure 2b where it can be seen that they approach the expected equilibrium concentrations.

- A second-order irreversible reaction: $A + B \overset{k}{\rightarrow} C$

If $[A]_0 = [B]_0$ then:

$$\frac{1}{[A]} = kt + \frac{1}{[A]_0} \quad (6)$$

and so it is expect that a plot of 1/[A] vs time will have slope k and intercept $1/[A]_0$.

This model was implemented in STEPS with species A located in vesicle surfaces, and species B in the cytosol. The species interact through the second order reversible reaction with an initial count of 1000 each (100 molecules of species A for each of the 10 vesicles, and 1000 molecules of B in the cytosol). The reaction rate was $2\mu M^{-1}s^{-1}$ and the simulation was run for 1000 iterations. Figure 4c compares output from STEPS to the analytical solution of equation 6.

The above is for the special case where the initial (and resulting) concentration of A and B are equal to each other, but if that is not the case then:

$$ln\left(\frac{[B]}{[A]}\right) = ln\left(\frac{[B]_0}{[A]_0}\right) - ([A]_0 - [B]_0)kt \quad (7)$$

For this test, the vesicle surface species, A, was given a starting count of 500 (50 per vesicle) and the cytosolic species, B, was given a starting count of 250 and they were allowed to react with each other at a rate of $0.5\mu M^{-1}s^{-1}$, with the simulation run for 1000 iterations. Figure 4d shows the output of the STEPS simulation compared to the analytical solution of equation 7.

### 10) Vesicle binding reactions

To validate the binding interactions between vesicles (and formation of 'link' species), we created a model that followed the kinetics of equation 6 because only one type of species was involved in the binding interaction. 100 small vesicles of 10 nm diameter and with 1 binding species in each vesicle surface were injected into the cytosolic volume of a spherical mesh of 0.5μM diameter. The binding interaction then took place with a reaction rate of $2\mu M^{-1}s^{-1}$ and

100 iterations of the simulation were run. Figure 4e shows the output of this model to the analytical solution of equation 6.

### 11) Raft diffusion

The 2D solution to the diffusion equation from a point source is:

$$C(r,t) = \frac{N}{4\pi Dt} e^{\left(-r^2/_{4Dt}\right)} \quad (8)$$

where C is the density (i.e. the number of molecules per unit area), N is the total number of molecules, D is the diffusion rate, and r is the distance from source.

The simplest test of raft diffusion in STEPS is to test conditions that approximate the point source by injecting rafts initially in an area that is small compared to the range of diffusion. We created a mesh of a flat circular disk shape with a radius of 10μm and injected rafts of 10nm radius close to the center of one of the circular faces, then allowed them to diffuse at a rate of 10μm$^2$/s. Overall, 100 rafts were injected. The density of rafts on the circular face was then compared to the analytical solution of equation 8 at times 30ms, 50ms, 70ms, 90ms and 110ms, as shown in Figure 5a, as the mean of 1000 iterations of the simulation. Boundary effects were negligible because very few rafts reached the 10μm circular edge at these times.

### 12) Raft molecule reactions

Reactions involving raft surface molecules were tested quite similarly as for section 9, with the difference that the reactants were situated on rafts undergoing diffusion on the mesh surface, and with some variation in parameters. 10 rafts of 100 nm diameter were placed randomly on the surface of a mesh of a 0.5μm diameter sphere and allowed to diffuse at a rate of 1μm$^2$/s.

Firstly, first order irreversible reactions were tested against equations 1 and 2, as shown in Figure 5b. This model was initialized with 50 molecules (5 per raft) which decayed by a rate of 5/s. Figure 5b displays the output of this model against equations 1 and 2 for 500 simulation iterations. For the first order reversible interactions, 1000 molecules were initialised per raft with a forward rate constant of 100/s and backward rate constant of 20/s. Figure 5c shows the mean output of this model against equations 4 and 5 for the equilibrium concentrations, over 10 simulation iterations.

Rafts themselves in STEPS can undergo special types of first order reactions that can generate or degrade rafts and can be tested in a production and degradation model. For simultaneous degradation and production such as: $A \overset{k_1}{\rightarrow} \emptyset$ , $\emptyset \overset{k_2}{\rightarrow} A$
the stationary distribution is:

$$\Phi(n) = \frac{1}{n!}\left(\frac{k_2}{k_1}\right)^n e^{-\left(k_2/_{k_1}\right)} \quad (9)$$

We implemented a model that could be compared to equation 9, with a raft generation reaction representing production and a raft dissociation reaction representing degradation. Rafts were generated from a species signature of just 1 molecule, with the number of these molecules in the mesh surface clamped at 1 throughout the simulation. This gave steady production of rafts, at a rate of 10/s. Degradation of rafts also occurred at a species signature of 1 molecule in the raft, at a rate of 1/s. The resulting number of rafts is then expected to follow equation 9. Figure 5d compares the output of the STEPS simulation to the analytical solution for a long simulation time of 100,000s (with samples taken every 0.1s) in order to negate any sampling error.

For the interactions between rafts and cytosolic species, we implemented second order reactions that could be comparted to equations 6 and 7. In the first test, 1000 raft surface species (100 per raft) could react with 1000 cytosolic species at a rate of $2\mu M^{-1}s^{-1}$. Figure 5e compares the output of the STEPS simulation (mean of 500 runs) to equation 6 for this model. In the second test, 500 raft surface species (50 per raft) reacted with 250 cytosolic species at a rate of $1\mu M^{-1}s^{-1}$, with the output from STEPS (mean of 500 runs) compared to equation 7, as shown in Figure 5f.

### 13) Vesicle occupancy effect on distribution of regular SSA species

Because the presence of vesicles can affect the diffusion of regular SSA species by a volume occupancy effect (as detailed in Methods) we tested the distribution of cytosolic species in a model with 10% occupancy by vesicles. We created a mesh of a spherical volume of a radius 10μm but of just 49 tetrahedrons for analytical purposes. Vesicles of diameter 50nm were inserted into the volume, comprising 4728 vesicles in total to give 10% occupancy, and they diffused at a rate of $1\mu m^2/s$. 1000 molecules were also injected into the volume and diffused at a rate of $10\mu m^2/s$. The simulation was run for 100s with the population of species recorded per tetrahedron at 1ms intervals. The distribution per tetrahedron is then expected to follow a binomial distribution, based on the reduced volume (the volume of the tetrahedron that is not occupied by vesicles). For this reason, the reduced volume was recorded per tetrahedron also at 1ms intervals.

Figure S3 shows the resulting distribution of molecules for all 49 tetrahedrons recorded from STEPS alongside the expected binomial distribution based on the mean reduced volume. The reduced volume can be seen in Figure S4 (a) for each tetrahedron. Figure S4 (b) summarises the recorded data shown in Figure S3 by displaying the error in the binomial fit compared to the expected binomial for all 49 tetrahedrons, showing low error across all tetrahedrons.

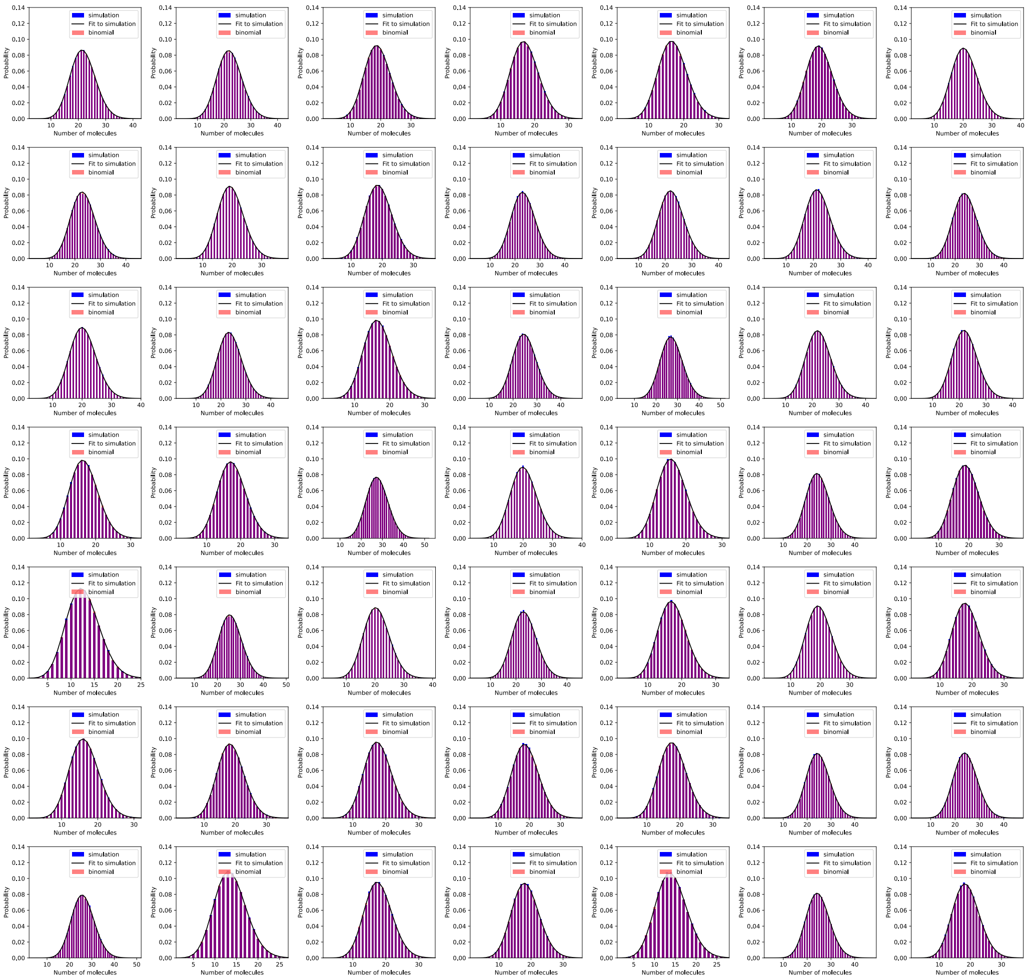


*Figure S3. In a model with 10% volume occupancy by vesicles, distribution of molecules over time in a STEPS simulation (blue bars) and binomial fit to that data (black line) compared to the expected binomial distribution (red bars) for all 49 tetrahedrons in the simulation volume.*

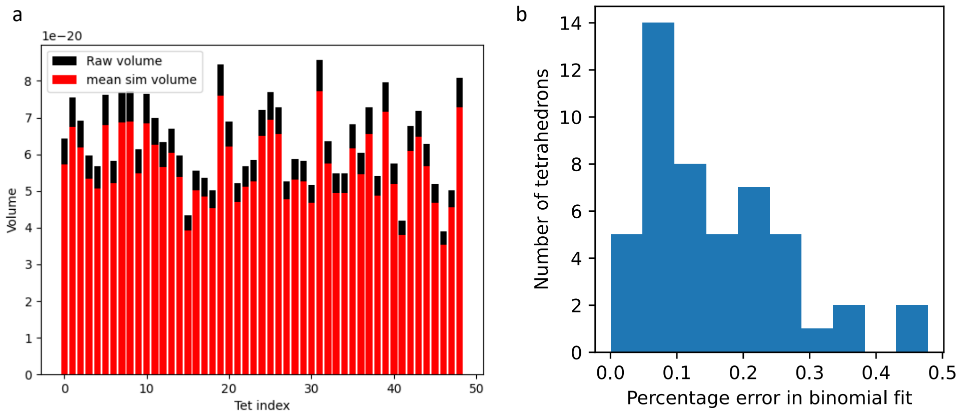


*Figure S4. a) the recorded reduced volumes of tetrahedrons (red bars, m³) in the 10% volume vesicle occupancy model compared to the unoccupied volume (black bars) showing the occupancy by vesicles as the difference. b) summary of the data shown in Figure S3 as the percentage error in binomial fit to the data compared to the expected binomial.*
